# GeneSetR: A web server for gene set analysis based on genome-wide Perturb-Seq data

**DOI:** 10.1101/2023.09.18.558211

**Authors:** Omer F. Kuzu, Fahri Saatcioglu

## Abstract

Identification of genotype-phenotype relationships is of utmost importance and a core effort in biology. Recent developments in efficient and precise gene targeting approaches coupled to omics methods have significantly improved deciphering of molecular interactions and relationships. However, many single gene perturbations can affect the expression of hundreds of other genes and analysis of the resulting omics-derived gene lists currently remains a significant challenge. Here we present Perturb-Seq based Gene Set Analyzer (GeneSetR), a user-friendly web-server that can analyze user-defined gene lists based on the data from a recently published genome-wide Perturb-Seq study, which targeted 9,866 genes with 11,258 sgRNAs in the K562 cell line. Through this tool, users can cluster gene lists following dimensionality reduction by various algorithms, undertake network analysis from RNA sequencing data, identify key nodes among the submitted genes, perform gene signature analyses, and generate heatmaps based on perturbation or gene expression data. GeneSetR enables researchers to readily identify gene clusters associated with specific phenotypes or biological processes, providing insights into the potential functional roles of these clusters and the role of single genes in them. With robust analysis capabilities, GeneSetR is a powerful resource to facilitate the exploration of genotype-phenotype relationships.

## INTRODUCTION

Understanding the relationship between genetic perturbations and the resulting phenotypic outcomes is a key challenge in molecular and cell biology. Historically, the analysis of gene lists derived from omics experiments was used to identify and infer the biological processes, pathways, and molecular functions that might be perturbed or affected by the specific experimental condition under study. Many tools such as Ingenuity Pathway Analysis (IPA), WebGestalt, DAVID, GSEA, g:Profiler and KEGG were developed for the analysis of omics-derived gene lists (1–6). However, these tools have substantial limitations. A significant issue is their inherent bias towards well-documented pathways. This bias, originating from dependence on the published literature and predefined gene lists, can skew data representation, making it less reflective of the entire biological landscape.

Furthermore, human curated databases are inherently limited in scope, while those developed using computational approaches often suffer from low reliability due to high false-positive rates. Commercially available tools such as IPA employ proprietary databases that are not accessible to the public. This restriction raises concerns about the comprehensiveness of the data and creates a financial barrier for many laboratories due to the cost of these tools.

Moreover, databases used by these tools aggregate data from varied sources such as different cell lines, tissues, and organisms, including those from less reliable sources. Given that many pathways behave differently across various systems and under diverse conditions, such aggregated data may not yield accurate or contextually relevant insights. Compounding this issue, many of these tools lack the capability to construct novel networks; instead, they carry out statistical analyses to determine if particular gene classes are overrepresented in the given gene list (such as GSEA, DAVID), or, at best, they overlay the submitted gene lists onto pre-established canonical pathways (for example, KEGG, Reactome).

Recent technological advances, e.g. genome-wide CRISPR screening studies coupled to affordable high throughput sequencing, revolutionized the identification of novel gene-phenotype relationships. In particular, the development of Perturb-Seq (also known as CRISP-seq and CROP-seq), a powerful method for determining the consequences of large number of genetic manipulations on gene expression profiles at the single-cell level has emerged as a tool that can shift paradigms in our understanding of cellular processes. By allowing for a detailed investigation of individual cellular responses, Perturb-Seq uncovers layers of complexity in genotype-phenotype relationships that were previously inaccessible (7–9).

Since its initial development in 2016, Perturb-Seq has been used to investigate a diverse range of biological processes. Initial studies utilizing Perturb-Seq provided novel insights into key biological processes, including endoplasmic reticulum (ER) stress signaling, regulatory pathways related to innate immunity, and cell cycle regulation (7–9). More recently, various additional biological processes have been probed using Perturb-Seq, such as the study of 35 autism spectrum disorder-related genes in vivo (10), the consequences of oncogene or tumor suppressor gene mutations in cancer (11), mechanisms of T cell exhaustion (12), and host-virus interactions during cytomegalovirus infection (13), among many others. In these studies, Perturb-Seq was limited to targeting only a few dozen genes, primarily due to the prohibitive costs of the next-generation sequencing required per experiment. The development of a novel high-throughput sequencing approach (14) facilitated the first Genome-Wide Perturb-Seq (GWPS) study, which analyzed the transcriptional consequences of 9866 genetic perturbations in the K562 chronic myeloid leukemia cell line (15). While GWPS is an extremely powerful technique and can be of immense value, the sheer volume of data generated presents a significant challenge for a typical user to extract meaningful information from it. Importantly, the GWPS data can be exploited to generate novel methods to analyze gene lists, which may overcome many of the limitations of the traditional tools. Recognizing this potential and the pressing need to maximize the utility of GWPS datasets, we have designed and developed GeneSetR, a Perturb-Seq based Gene Set Analyzer.

GeneSetR, available at https://genesetr.uio.no, sidesteps many of the limitations of other available gene list analysis tools by anchoring its analysis on specified Perturb-Seq experiments. This approach mitigates reliance on broad, lowly curated databases that are likely contaminated. By focusing on a more specific and highly controlled dataset, GeneSetR offers a novel tool for gene list analysis.

## METHODS

### Datasource

GeneSetR primarily uses the GWPS dataset that has recently been published by Replogle et al. (15), but it will incorporate data from similar studies as they become available. In this study, a GWPS was performed on K562 leukemia cells; after filtering for various quality metrics, scRNA-seq data was collected from more than 2.5 million cells, expressing 11,258 sgRNAs that target 9,866 distinct human genes. On average, more than 100 cells were detected per perturbation and more than 85% of the sgRNAs successfully decreased the expression of their respective targets by greater than 50%. Two additional but smaller Perturb-Seq experiments were also performed, targeting 2,057 common essential genes in K562 and RPE1 cells (a non-cancerous hTERT-immortalized retinal pigment epithelial cell line). Although these two small datasets can also be analyzed in GeneSetR, due to their smaller size, they have limited utility at present.

The GWPS dataset is publicly available through a web page (https://gwps.wi.mit.edu) and is a valuable resource for researchers studying gene expression patterns. However, the limited set of tools that are available at this site is insufficient for more in-depth analyses. The available tools include mapping the location of a single gene on the embedding of a subset of perturbations with strong transcriptional phenotypes, visualizing the top 30 most up- or down-regulated genes for a selected perturbation, or assessing the top 10 most similar perturbations to the selected one, which are limited in scope. Moreover, some of these tools have limitations that can affect the accuracy of the results. For instance, the similarity assessment in the perturbation correlation is based on the Pearson correlation method that is sensitive to outliers; thus, an extreme gene expression value may interfere with the correlation of many perturbations, leading to inaccurate results. For example, PRG2, encoding proteoglycan 2, is highly upregulated by many perturbations, including PIK3C3. Because of its outlier expression, all genes that lead to extreme upregulation of PRG2 are annotated as if they are very highly correlated with each other, which in reality is not the case. Furthermore, heatmap generator from user-supplied lists of genes and perturbations is limited in scope, e.g. clustering on rows or columns is not possible, making it challenging to identify patterns in the data. We therefore developed GeneSetR to address these limitations and to significantly improve the accessibility of the invaluable GWPS dataset for the research community.

### GeneSetR modules

The GeneSetR encompasses a diverse array of modules, each specifically designed to provide powerful functionalities utilizing the high-dimensional data derived from Perturb-Seq studies. It contains modules for Dimensionality Reduction and Clustering, Correlation, Pathway Exploration, Gene Expression Analysis, Gene Signature Analysis, and Heatmap generation. Each module is designed to provide distinct components that aid in uncovering meaningful insights from the complex GWPS data. Below is a summary of each module.

### Dimensionality Reduction and Clustering Module

In the analysis of large datasets, including those generated in Perturb-Seq studies, there are often a large number of features, such as identified genes and specific perturbations. This high dimensionality can make it challenging to visualize and model the data, and can also lead to a significant increase in computational burden for downstream analyses such as clustering. To address these issues, dimensionality reduction (DR) techniques are commonly used to reduce the number of features while preserving as much of the original information as possible, by replacing redundant features with new ones that are derived from the combination of original ones.

DR algorithms can be broadly categorized into two types based on the underlying mathematical and computational approaches (for reviews, see (16–18)). Linear DR algorithms seek to find a lower-dimensional representation of the data by creating linear combinations of the original features. These combinations, or principal components, are chosen to maximize the variance of the data and minimize the redundancy between the original features. In contrast, non-linear DR algorithms aim to preserve non-linear relationships between features in the lower-dimensional space. They achieve this by using more complex mathematical functions to map the original features onto a lower-dimensional space. Linear DR methods, such as Principal Component Analysis (PCA), are popular because they are computationally efficient and can easily be interpreted. Non-linear DR methods are more computationally demanding, but can capture more complex relationships between features. Thus, depending on the goal, complexity of the data, and resource availability, it may be desirable to use different DR methods or their combinations.

In GeneSetR, users can apply the four most common DR techniques that are used in scRNA-seq data analyses: PCA, t-Distributed Stochastic Neighbor Embedding (tSNE), Uniform Manifold Approximation and Projection (UMAP), and Multidimensional Scaling (MDE). In the PCA module of GeneSetR, the only parameter that users are expected to set is the threshold variance level that is used to automatically determine the minimum number of components required to explain that much variance.

On the other hand, tSNE, UMAP, and MDE are non-linear DR techniques. They work by modeling the probability distribution of each point in the dataset and then mapping the points to a lower dimensional space in a way that preserves the local structure of the data. In contrast to PCA, these algorithms provide more accurate representation of the underlying data structure. However, they are computationally more expensive, and may take significantly longer to calculate, or fail if the input gene list is too long. In such cases, users are encouraged to perform PCA prior to applying these DR algorithms, particularly before tSNE or MDE. Each of these algorithms have several parameters, and unfortunately, there is no one-size-fits-all solution. The optimal parameter values may vary depending on the submitted gene list, and should be tuned to achieve a better visualization of the high-dimensional data. GeneSetR provides tool tips on these parameters to guide users in dimensionality reduction optimization.

Following dimensionality reduction, GeneSetR employs Hierarchical Density-Based Spatial Clustering of Applications with Noise (HDBSCAN) algorithm to cluster the data (19). We use this clustering method as it has several advantages over other commonly used clustering algorithms. First, it is a density based algorithm and hence can produce clusters of varying shapes, sizes, and densities without requiring any prior knowledge about the data. This is particularly useful when dealing with high-dimensional data, where clusters may have complex shapes. Second, HDBSCAN does not require the user to specify the number of clusters, making it more intuitive and easier to use than other methods that rely on this parameter. However, despite its advantages, there are still a few parameters (e.g. minimum cluster size, minimum number of samples per cluster, and clustering metric) that users need to tune for finding the optimal number of clusters in their data. Adjusting these parameters can help users achieve the best possible clustering results for their given gene list.

If HDBSCAN clustering identifies clusters for the input gene list, GeneSetR offers an intuitive tool for conducting GSEA on the identified clusters. Leveraging the power of the EnrichR API, GeneSetR utilizes the Gene Ontology Biological Processes (GO-BP) library as the default annotation database, with the option to expand to include other available annotation libraries in the EnrichR tool (20). The enrichment results are visually presented through interactive bubble charts and can be further explored in a downloadable table format.

### Correlation Module

This module enables users to create heatmaps based on perturbation or gene expression data. Users can select between two correlation methods, Pearson and Spearman, and the analysis can be performed using all of the GWPS data set or a subset of the data. For example, users can enter a perturbation list and submit a gene list to limit the correlation calculation to only a certain pathway or a family of genes included in the gene list. If the gene list is empty, the calculation will be based on all available data.

To visualize the results, the Correlation Module uses a modified version of the Interactive Cluster Heatmap library (Inchlib) (21), which can perform hierarchical clustering on both rows and columns. Although users are free to apply different clustering settings (such as distance metrics) to rows and columns, the resulting heatmap would reveal gene or perturbation clusters as distinct rectangular blocks along the diagonal, if the settings are kept the same **(Figure 1)**. One of the advantages of the GeneSetR Correlation Module is its ability to provide a comprehensive overview of the relationship between gene expression and perturbations. This can help users to identify patterns in their data that may not be immediately apparent. Additionally, the ability to limit the correlation calculation to specific pathways or gene families can provide more targeted insights into gene regulatory mechanisms.

**Figure 1.**
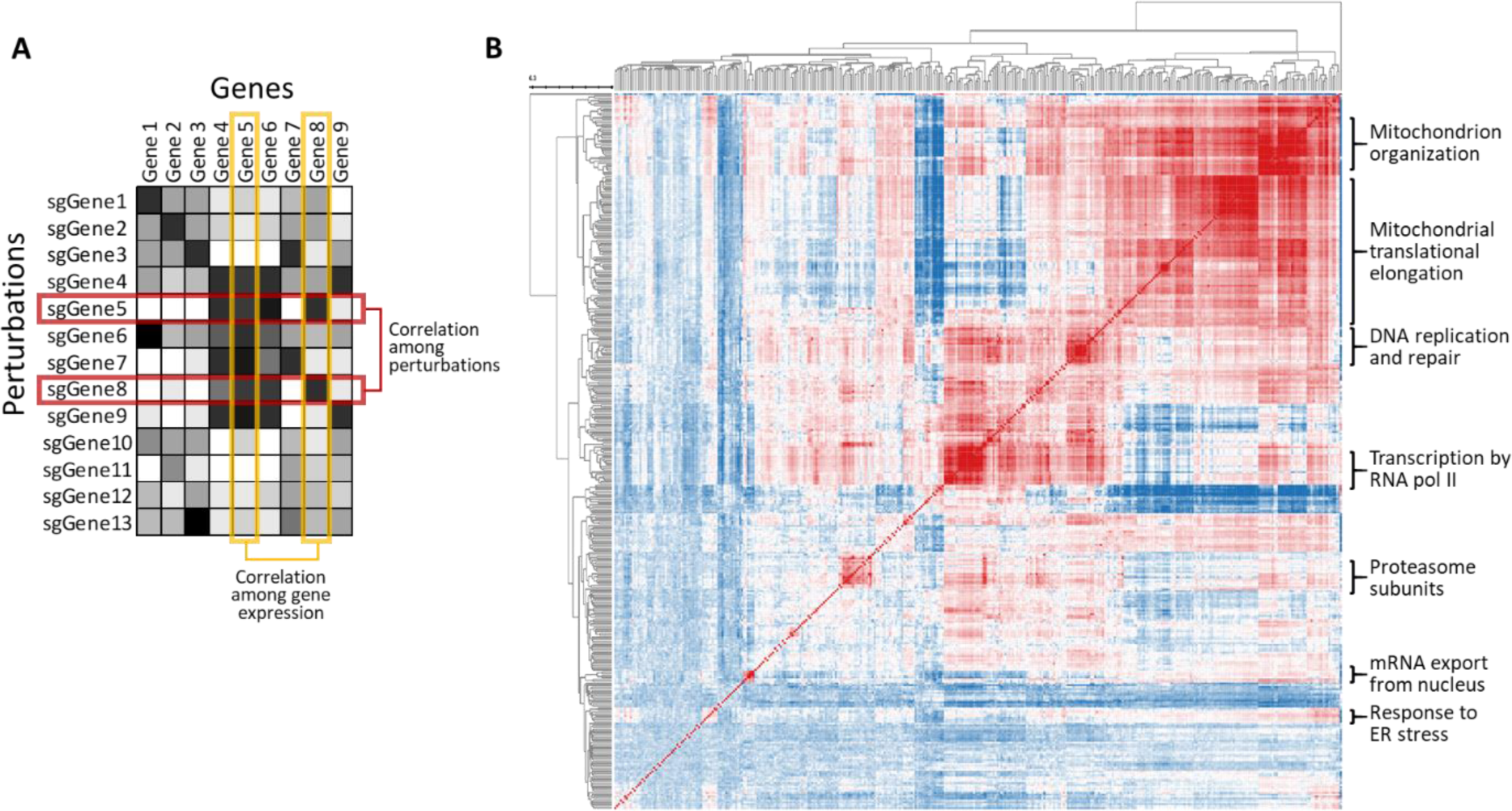
Correlation module generates functional gene clusters. In this module it is possible to generate correlation heatmaps based on correlation of perturbations or gene expression. **A.** The figure illustrates a hypothetical example from Perturb-Seq data where 13 perturbations are considered. Analysis can be undertaken both at the perturbation level (correlating rows) and the gene expression level (correlating columns). Notably, in this example, a marked correlation is evident between the perturbations of Gene 5 and Gene 8. However, the responses of these genes to various perturbations (i.e., the correlation among their expressions) do not seem to be correlated. **B.** A Pearson correlation heatmap, derived from perturbation correlations, was generated for 384 genes that were identified in a CRISPR screen (see Results for the details of this gene list). Note that distinct rectangular blocks along the diagonal reveal functional clusters (annotated at the right).

### Pathway Explorer Module

This is one of the most useful features of GeneSetR. It utilizes GWPS data to generate pathways among submitted genes, making it particularly valuable for analyzing results from RNA-seq experiments **(Figure 2)**. To generate pathways, the Pathway Explorer Module requires a list of down-regulated genes (and optionally upregulated ones) derived from a differential gene/protein expression analysis experiment (e.g., RNA-Seq or proteomics). The algorithm examines each down-regulated gene on the list and determines, based on a z-score threshold chosen by the user, as to which genes are up- or down-regulated following the perturbation of that particular gene in the GWPS experiment. If the genes that are up- or down-regulated also appear among the submitted list of up- or down-regulated genes, an edge connecting the perturbed gene and the target is created. Depending on the “depth” setting, this process is performed either solely on the submitted down-regulated genes, or also on genes that are down-regulated upon perturbation of these genes in the GWPS data **(Figure 2)**. Moreover, the Pathway Explorer Module can annotate created pathways with the perturbation correlation data from the GWPS experiment, or with the protein-protein interaction data from the BioGRID database (https://thebiogrid.org). This information is then shown as additional edges between the nodes of the pathway.

**Figure 2.**
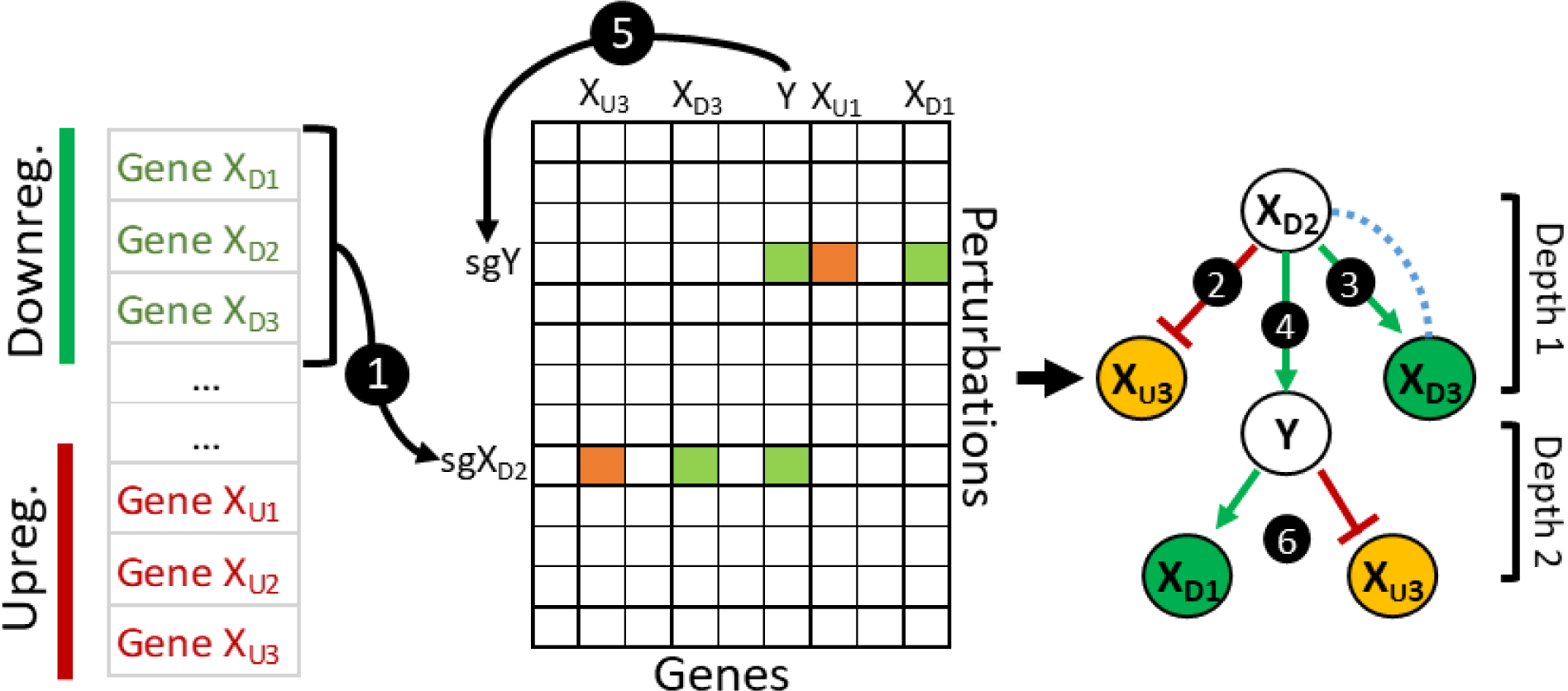
Pathway Explorer Module generates pathways among the submitted genes. The down-regulated genes are scanned and the ones that are targeted in the Perturb-Seq data are examined for genes that are regulated upon their knockdown (1). If a down-regulated or up-regulated gene exists among the submitted gene lists that are regulated in the same direction, an edge between those are created (2–3). If the depth setting is set to two, another round of scanning is performed including intermediate genes (such as Y) (4 and 5).

In addition to its utility in analyzing RNA-seq data, the Pathway Explorer Module is also instrumental in identifying key nodes among any set of submitted genes. For example, as demonstrated below in Section 4.2, when a gene list obtained from a genome-wide CRISPR screen is used, GeneSetR can unveil the interactions among the submitted genes, revealing the key nodes that mediate the observed phenotype. The resulting network is visually presented using the Apache ECharts graphing library, with node sizes adjusted based on the count of their interaction partners, and node opacity determined by the knockdown efficiency of the gene in the GWPS data. Similarly, the width and color opacity of the links highlight the effect size, providing comprehensive, intuitive and visually appealing insights into gene-gene interactions.

### Gene Expression Analysis Module

GeneSetR also features a Gene Expression Analysis Module where users can interrogate a single gene of interest (GOI) for its potential function. This module facilitates: (1) identification of genes that are up- or down-regulated upon targeting the GOI (downstream targets), (2) pinpointing perturbations that can induce or repress expression of the GOI (upstream regulators), (3) discovering perturbations that correlate with the targeting of the GOI, and (4) identification of genes that display similar expression responses as the GOI upon perturbation of other genes.

In the Gene Expression Analysis Module, users can create networks centered on a GOI. The initial step involves identifying upstream regulators of the selected gene based on a user-defined z-score threshold. Provided that the GOI was successfully targeted in the GWPS data (it is worth noting that only approximately half of the known genes were targeted, but this will change as more GWPS data become available), the network is expanded to include downstream targets (genes that are influenced by the knockdown of GOI).

Since upstream regulators of the GOI are expected to also regulate its downstream targets, links between upstream regulators and downstream targets are examined. Only those links that are consistent with expectations (e.g., an upstream positive regulator (UPR) would positively regulate a downstream positively regulated (DPR) target and negatively regulate a downstream negatively regulated (DNR) target, while an upstream negative regulator (UNR) would do the opposite - see **Figure 3a**) are incorporated into the network. Similarly, to enhance the reliability of the network, intra- and inter-group interactions among the UPR, UNR, DPR and DNR genes are also examined; thus, in total up to 16 different logical interaction sets are evaluated and added to the network where applicable. Moreover, correlations between perturbations of the nodes are highlighted when they align with the flow of the pathway. These logical links around the GOI would assist users in identifying novel and reliable links and offer a comprehensive view of the regulatory pathway(s) surrounding the GOI **(Figure 3b)**. In addition to allowing user-friendly visualization of the generated network, this module also provides users with the ability to view and download the underlying data as tables.

**Figure 3.**
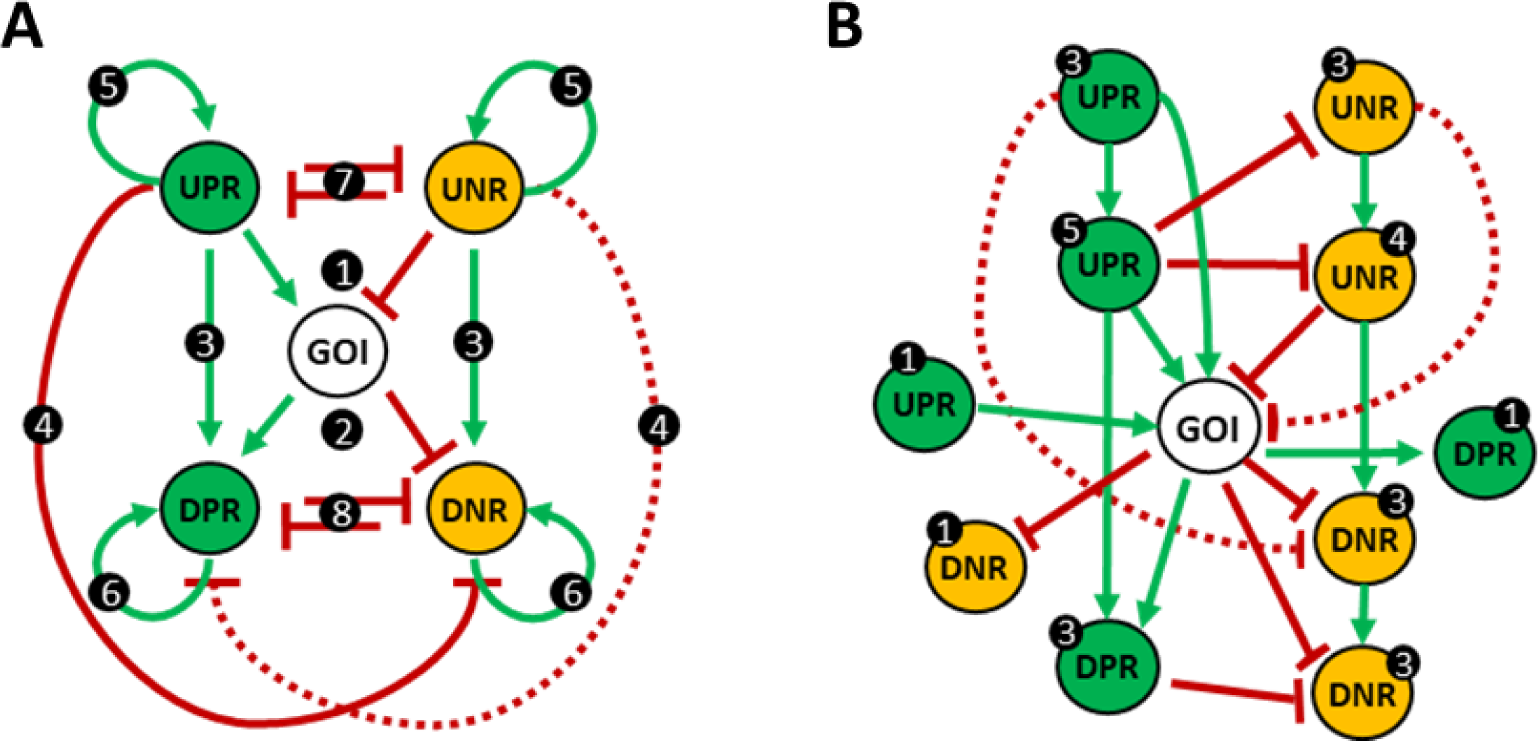
The Gene Expression Module generates a network surrounding a gene of interest. a. If the **G**ene **O**f **I**nterest (GOI) is detected in the Perturb-Seq dataset, upstream genes (perturbations) that regulate the GOI are identified (**U**pstream **P**ositive/**N**egative **R**egulators-UPR/UNR) (1). Next, if a sgRNA exists for the GOI, genes that are up-regulated (**D**ownstream **P**ositively **R**egulated-DPR) or down-regulated (**D**ownstream **N**egatively **R**egulated-DNR) upon knockdown of the GOI are added to the network (2). To strengthen the network, all logical links among upstream regulators and/or downstream targets are also appended to the network (3–8). **b.** An example network generated around a putative GOI. The number of neighbors for each gene is highlighted in black circles. These numbers would indicate the strength or reliability of that particular gene’s involvement in the network.

The Gene Expression Analysis Module often generates a very dense network that necessitates filtering to focus on the most relevant connections. To address this, the module offers a range of filters to streamline the visualization, thereby facilitating a more accessible interpretation of the underlying biological processes. Key filters include options for setting thresholds for interaction strengths (e.g. knockdown z-scores or correlation coefficients), including or excluding various types of interactions (e.g. UPR to DPR), displaying only genes with a minimum number of interactions, and excluding isolated genes (those without any links to other genes).

Single cell RNA-Seq experiments are prone to different types of “noise”, or sources of variation, that can introduce bias into the results. The Gene Expression Analysis Module provides essential filters to mitigate two distinct sources of noise in the generated networks. The first type of noise is related to perturbations that influence overall mRNA levels. The second type of noise comes from the abnormal expression of specific genes in response to perturbations, either due to inherent biological traits (like high or variable expression) or technical biases in RNA-sequencing (like the overrepresentation of more abundant RNAs). For a comprehensive analysis of noise, please refer to the ‘Noise Filtering’ section provided in the Supplementary Information. An in-built filtering tool within the module allows for the exclusion of the “blacklisted” genes from the network. It is important to note, however, that while noise filtering options can enhance the clarity and reliability of the results, they should be used judiciously, keeping in mind the biological context and the specific goals of the analysis.

### Gene Signature Analysis Module

GeneSetR has the capability to perform gene signature analyses, which can identify genes that induce specific phenotypes upon their knockdown. Users can provide mathematical expressions (summation or subtraction), which GeneSetR applies to z-score normalized data and returns the list of perturbations with resulting average values. During the calculations, members of the gene signature are excluded from the formula when calculating the score for that particular gene, as the CRISPRi-mediated knockdown of the gene would overshadow its score (e.g., if a gene signature contains the AKT1 gene, during the calculation of a score for sgAKT1, AKT1 is excluded from the signature). Perturbations that significantly alter the gene signature are identified by calculating z-scores, using a 0.2% trimmed mean and standard deviation. This approach reduces the false negative rate caused by outlier values. With this module, researchers can efficiently and accurately identify set of genes that are responsible for a specific phenotypic change, as demonstrated below in the “Pathway Explorer analysis” subsection in Results.

### Heatmap Module

This module enables users to create visually informative heatmaps from the GWPS data. With the ability to input a list of genes and perturbations, this module renders gene expression data as a heatmap using the Inchlib library (21). In this setup, users have the flexibility to perform clustering on both rows and columns. The ability to customize clustering parameters allows users to tailor the analysis to their specific needs. The heatmap provides an intuitive visual representation of gene expression levels, allowing users to identify trends and patterns in the data at a glance. Users can then zoom into the clusters of interest, create gene lists from the clusters, and perform GSEA with a single click. With its visually intuitive interface, the Heatmap Module facilitates deeper exploration of the GWPS data and enables better understanding of gene expression dynamics in response to perturbations.

### Implementation

GeneSetr is developed on a backend powered by Flask and Python, with Celery handling task management. The tool utilizes the Redis database to transiently store results from Celery tasks. On the frontend, developed with React, the software employs the Echarts graphics library for data visualization, and the Inchlib library is incorporated for specialized heatmap and correlation modules. GeneSetr integrates the EnrichR API to perform geneset enrichment analyses. The platform does not necessitate user login and ensures data privacy by locally storing genelists and enrichment results using the Web Storage API.

## RESULTS

To illustrate its capabilities, we utilized GeneSetR to analyze a gene list acquired from a recent CRISPRi screen in K562 cells that expressed a mCherry reporter driven by a 5X unfolded protein response element (7). In this study, to identify genes that would lead to endoplasmic reticulum (ER) stress upon their knockdown, the cells were transduced with a genome-wide CRISPR library targeting 18,905 genes (on the average one sgRNA per cell), and then grown for 8 days. Flow cytometry was performed to quantitate the frequencies of sgRNAs contained in the top and bottom third of mCherry expression, which identified 397 hit genes with high mCherry expression suggesting activation of unfolded protein response signaling.

As noted above, GWPS data targets only about 50% of the human gene set; thus, GeneSetR automatically checks the availability of the entered genes and recognizes gene symbol aliases to prompt users for alternative gene symbols to ensure that users would be able to easily map their gene list. In the data set from the ER stress study described above, 13 of the 397 genes were not targeted in the GWPS study and therefore only the remaining 384 genes were used in the analysis below.

### Dimensiality Reduction and clustering analyses

First, we performed PCA analysis on these data limiting the threshold value for the amount of variance that needs to be explained to various values in the range of 50 to 100 percent. In PCA, as long as there are more than two principal components, altering this setting would not affect the displayed PCA plots, as increasing the number of principal components would not alter the first three components that are used to draw these plots. However, the number of calculated components can have a significant impact on the clustering of the data. With a 50% variance explanation threshold, 13 components were calculated which led to four clusters by the HDBScan algorithm. Increasing the threshold to 60% increased the component number to 27 resulting in identification of an additional cluster **(Figure 4)**. Further increases in the threshold gradually decreased the identified number of clusters. The decrease in clustering efficiency with increasing number of dimensionality is known as “the curse of dimensionality” where clustering algorithms become less efficient to identify meaningful clusters due to the sparsity and dissimilarity of samples in high-dimensional data and highlights the importance of careful selection of the appropriate settings for a given analysis (22).

**Figure 4.**
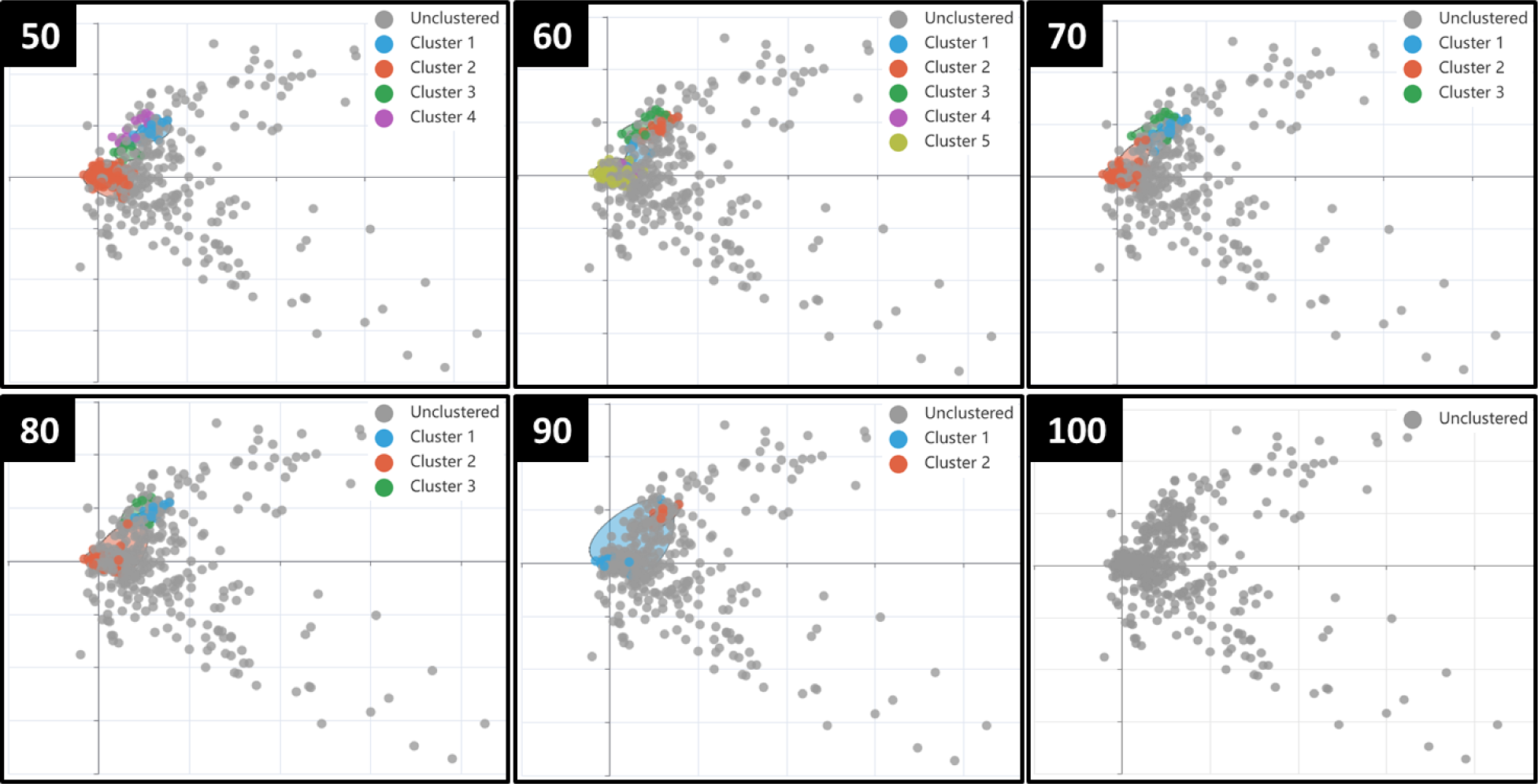
PCA analysis of perturbations on genes that induce ER stress. PCA analysis was performed for increasing values of variance explanation setting, as pointed out in the main text, at the range of 50 to 100 percent. Note that HDBSCAN based clustering analyses identified maximum five clusters at 60% EV, and identified number of clusters gradually decreased with increasing EV values.

GSEA using the GO-BP as the annotation database suggested that four of the five clusters that were identified in the PCA analysis (at 60% threshold level), were highly enriched for specific biological processes **(Table 1)**. For instance, the first cluster contained six genes five of which are involved in the aerobic electron transport chain. COQ2 was the only gene in this cluster that was not labeled as a gene involved in the aerobic electron transport chain, despite coding for an enzyme involved in the biosynthesis of ubiquinone (CoQ), a redox carrier in the mitochondrial respiratory chain (23). The second cluster contained 25 genes and according to the GO-BP database, 22 of them are involved in the mitochondrial translational elongation. PTCD1, DHX30, and TARS2 were the three remaining genes that were not labeled for the mitochondrial translational elongation activity despite multiple reports regarding their role in this process (24–26). Similarly, the third cluster contained 14 genes, in which 11 were also labeled for their involvement in mitochondrial translation. Three remaining genes in this cluster (TFB1M, METTL17, and ATP5ME) also encode mitochondrial proteins that are strongly involved in mitochondrial protein translation. The fourth cluster had 7 genes, and three of them were labeled for ER to cytosol transport. In this cluster, three (SLC39A7, YIPF5, TMED10) of the four remaining genes were ER resident proteins known to be involved in ER stress, whereas the last one, GMPPB, is involved in protein glycosylation, a process that is well-known to cause ER stress upon inhibition (27). Taken together, these examples clearly demonstrate that GeneSetR can cluster submitted gene lists better than the most complete annotation databases in an unbiased and literature-independent manner.

**Table 1.**
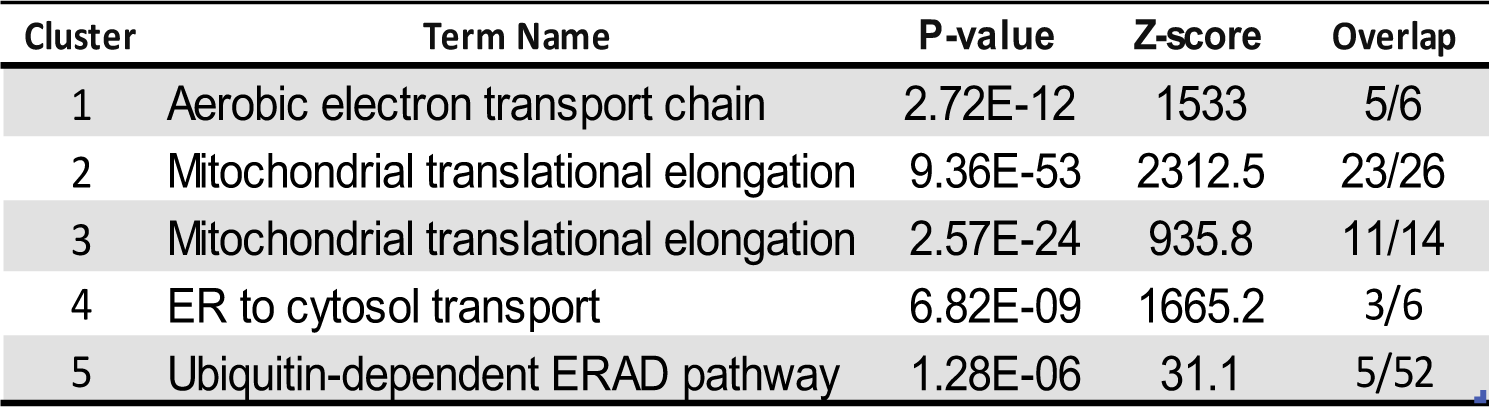
GSEA of clusters obtained upon PCA. GSEA was performed on clusters identified by PCA using Gene Ontology Biological Processes database (2021) through the EnrichR API. Note that four of five clusters are highly enriched for specific processes.

| Cluster | Term Name | P-value | Z-score | Overlap |
| --- | --- | --- | --- | --- |
| 1 | Aerobic electron transport chain | 2.72E-12 | 1533 | 5/6 |
| 2 | Mitochondrial translational elongation | 9.36E-53 | 2312.5 | 23/26 |
| 3 | Mitochondrial translational elongation | 2.57E-24 | 935.8 | 11/14 |
| 4 | ER to cytosol transport | 6.82E-09 | 1665.2 | 3/6 |
| 5 | Ubiquitin-dependent ERAD pathway | 1.28E-06 | 31.1 | 5/52 |

The number of components (NOC) setting holds critical importance for all DR algorithms. Therefore to systematically evaluate the optimal value for NOC setting, and to determine which DR algorithm would result in functionally relevant clusters, we conducted a benchmarking study using four DR algorithms, as well as some of their combinations (for details, see ‘Benchmarking of dimensionality reduction algorithms’ section provided in Supplementary Information). In our comparative analysis, PCA led to fewer clusters and peaked at a clustering efficiency score of 35 at 60% explained variance. UMAP demonstrated relatively consistent performance across varying NOC values, achieving a top score of 44. Maximum clustering score obtained with tSNE was 48, but it was very slow in computing and its efficiency was highly sensitive to the NOC and perplexity settings. MDE had difficulty handling high NOC values but reached the highest efficiency among individual algorithms at a specific NOC. Notably, the combined application of PCA with the non-linear algorithms, especially MDE, significantly improved clustering, with the PCA-MDE combination achieving the highest score of 80 at a NOC of 19, generating up to 23 clusters **(Table 2)**.

**Table 2.** Comparison of best benchmarking results. Summary of the best results obtained during benchmarking of the various dimensionality reduction algorithms. NOC = Number of Components value for each run.

| DR algorithms | NOC | Clustering Score | Cluster Count | Clustered Gene Count | Mapped Gene Count | Computation Time |
| --- | --- | --- | --- | --- | --- | --- |
| PCA-MDE | 21 | 80.0 | 22 | 391 | 118.4 | 1.56 |
| PCA-UMAP | 3 | 55.7 | 13.8 | 343.1 | 92.4 | 5.07 |
| MDE | 43 | 48.7 | 14.8 | 288 | 79.8 | 1.86 |
| tSNE (Perplexity 1) | 3 | 47.8 | 18 | 345.5 | 82.5 | 2.20 |
| UMAP | 3 | 43.1 | 9 to 12 | 339.2 | 70.9 | 5.15 |
| PCA-tSNE (Perplexity 10) | 10 | 36.6 | 9.2 | 134.6 | 48.4 | 11.33 |
| PCA | 60 | 34.5 | 5 | 104 | 43 | 0.36 |
| tSNE (Perplexity 10) | 3 | 11.2 | 4.6 | 121.5 | 20.3 | 3.29 |

### Pathway Explorer analysis

To highlight the utility of the Pathway Explorer Module, we conducted an analysis using the same set of 384 genes mentioned above, which induce ER stress upon their knockdown. When this gene list was run in the Pathway Explorer Module, several genes were found to be regulated by other genes in the list, suggesting that they could be potential key nodes in the induction of ER stress **(Figure 5)**. One notable example was RPL41, which encodes a peptide consisting of only 25 amino acids. With a z-score cutoff of 0.75, knockdown of 34 genes down-regulated RPL41 expression. Interestingly, RPL41 has been reported to induce rapid degradation of the ATF4 protein, which is a key transcription factor in the ER stress responses (28). Consistently, RPL41 knockdown effectively induced ATF4 levels, whereas treatment of cells with exogenously produced RPL41 suppressed it (28,29). These findings suggest that genes that positively regulate RPL41 expression may influence the reporter expression and could be inadvertently identified as hits in the CRISPR screen.

**Figure 5.**
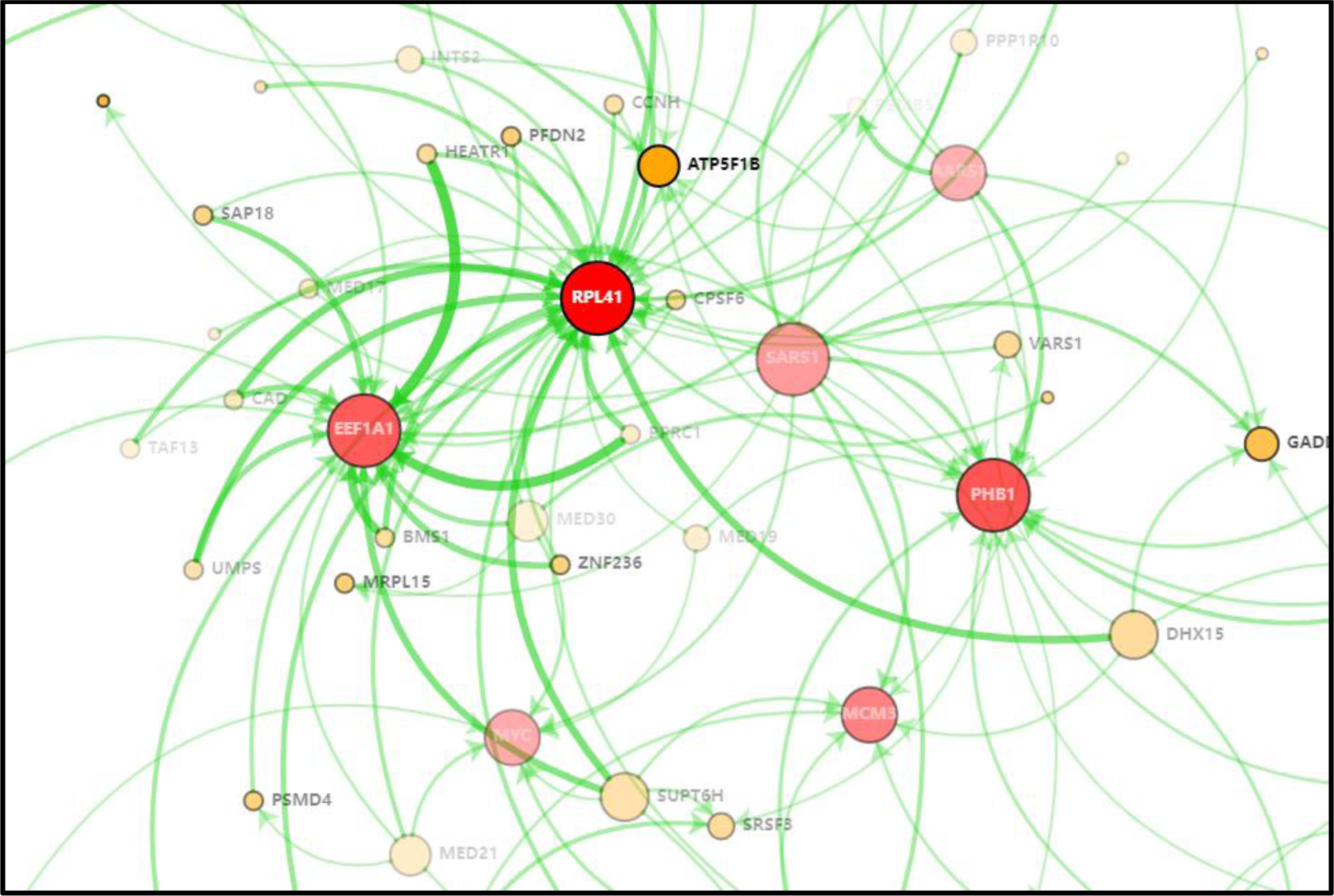
Pathway Explorer module identifies key genes among the queries. The figure displays the network generated by the Pathway Explorer module for 384 genes that induce ER stress upon their knockdown. It is noteworthy that certain genes, such as RPL41, EEF1A1, and PHB1, are down-regulated by numerous other genes and can be considered as key nodes within the network.

Another key node in the network was EEF1A1, with 20 neighbors at a 0.75 z-score cutoff. This gene encodes eEF1A1, one of the most abundant proteins in the cell, which is involved in the elongation phase of protein synthesis, facilitating the delivery of aminoacyl-tRNAs to the ribosome (30). However, eEF1A1 is not only a translation factor, but also exhibits chaperone-like activity and participates in cellular responses to various stresses, including heat-shock, proteasomal stress, and ER stress (30–32). Moreover, eEF1A1 has been reported to inhibit Gcn2-mediated eIF2α phosphorylation; thus, its decreased expression could also positively regulate ATF4 signaling (30).

A third key node in the generated network, PHB1, was down-regulated upon knockdown of 17 other genes in the list (at z-score cut off 0.75). This gene encodes a protein called prohibitin 1, which plays a key role in the maintenance of mitochondrial integrity (33). Accumulation of prohibitin 1 has been reported as a common cellular response to different stress stimuli in melanoma cells, where it protects cells from ER stress-induced cell death (34). In mice, deletion of Phb1 caused mitochondrial stress and ATF4 activation through the integrated stress response pathway (35). These findings also suggest that modulation of PHB1 expression could account for a portion of the identified CRISPR screening hits.

As shown in these examples, the Pathway Explorer Module serves as a powerful tool to identify key signaling nodes within gene lists of interest.

### Gene Regulation Analysis

As summarized above, the Gene Regulation Analysis Module enables a detailed dissection of regulatory networks linked to a single GOI. To demonstrate the capabilities of this module, we focused our investigation on the *ATF4* gene.

ATF4 is a transcription factor that acts as a downstream effector in integrated stress response signaling (36). Through its regulatory function, it influences a broad array of genes connected to cellular metabolism and stress response. This multifaceted role in maintaining cellular homeostasis is further complemented by its involvement in processes such as autophagy, apoptosis, and amino acid biosynthesis. The regulation of ATF4 expression and activity is a complex process, incorporating multiple stages of transcription, translation, and post-translational modifications, highlighting the anticipated complexity of the network associated with it. Given the complexity of its regulation, an expansive network encompassing ATF4 is anticipated. Corroborating this hypothesis, the Gene Regulation Analysis Module identified in excess of 40,500 links (at a z-score threshold of 0.4, excluding correlative links), in the absence of noise filtering. The introduction of four noise filters significantly reduced this massive network to 2492 links; although still extensive, this allowed for a more focused and manageable analysis of the regulation network **(Supp. Table 1 and 2)**.

While noise filters are instrumental for minimizing link volume, they can also inadvertently eliminate numerous genes (many ribosomal proteins and genes involved in mTOR signaling) known to be involved in ATF4 signaling, as outlined above, in the “Gene Expression Analysis Module” subsection of Methods. After removal of the noise and when the interactions are ranked based on the total number of neighboring links, the Gene Regulation Analysis Module demonstrates its efficacy in identifying the genes that either regulate or are regulated by ATF4 **(Figure 6 and Supp. Table 1)**. The top three genes — PHGDH, WARS1, and PSAT1 — are all well-established direct downstream targets of ATF4, with their connections supported by approximately 50 links each. The fourth gene, EIF2B1, a vital element in the regulation of ATF4 translation, is supported by 28 links, positioning it as an upstream negative regulator of ATF4. Among the top 20 genes, eight are tRNA synthetases, all of which have established associations with ATF4 signaling in the existing literature. This analysis further reinforces the module’s robust capability to accurately and effectively uncover the complex regulatory networks surrounding any GOI.

**Figure 6.**
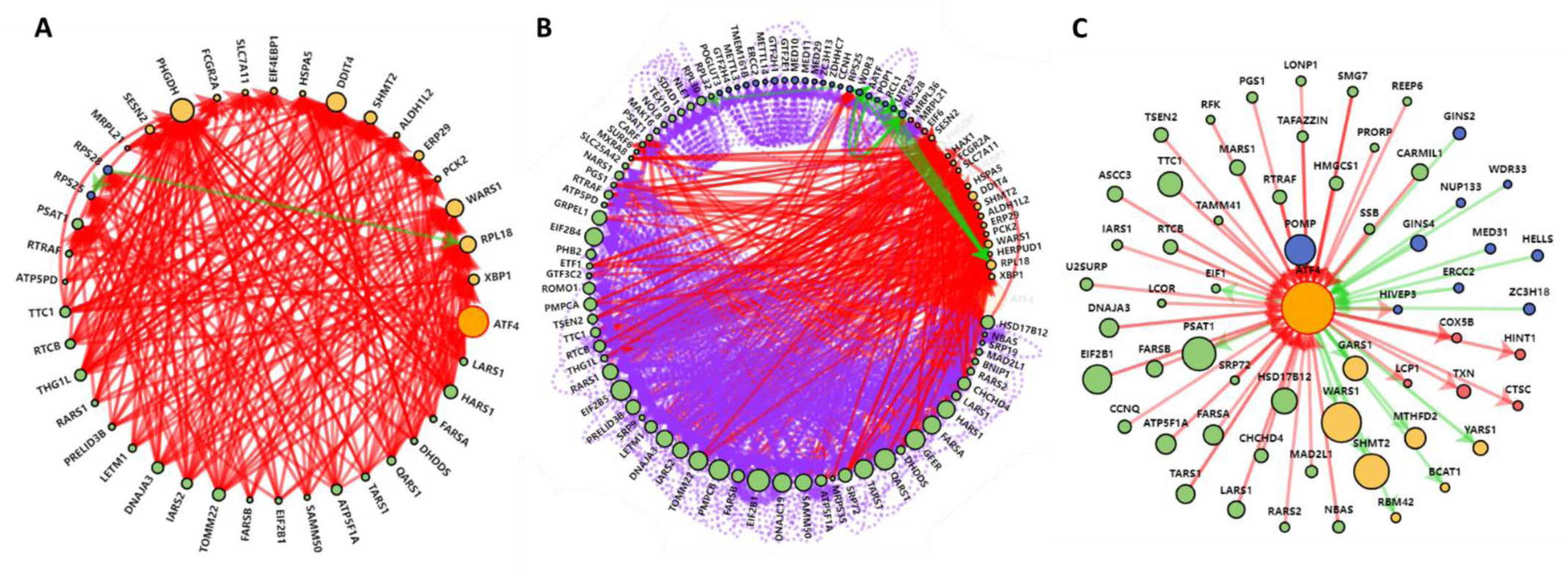
The Gene Expression Module generates a network around a given gene of interest. Network generated by the Gene Expression Analysis Module for the *ATF4* gene using the following thresholds: for a z-score of 0.7, at least 5 neighbors, excluding correlation (**a**), or including correlation with a 0.5 Spearman r cutoff (**b**). **c.** A simplified ATF4 network generated by the Gene Expression Analysis Module with the following thresholds: z-score of 0.4, at least 10 neighbors, excluding correlation (see Supp. Table 1 for details). Note that many well-known ATF4 upstream regulators (aminoacyl tRNA synthetases, genes that trigger integrated stress response upon knockdown) and downstream targets (PHGDH, SHMT2, SLC7A11, and EIF4EBP1) are included in the network.

### Gene Signature analysis

To evaluate the practical applicability of our gene signature analysis module, we used a gene signature that we have formulated based on up- or down-regulated genes in response to ER stress (7). This gene signature should be able to identify genetic perturbations that can activate the unfolded protein response.

In an unfiltered state, the gene signature identified 480 genes with a z-score > 2. Of these, 120 genes overlapped with the aforementioned set of 397 genes (384 of them available in GWPS dataset) **(Figure 7a)**. This gene list was previously utilized to illustrate the clustering analyses in the preceding section (7). As anticipated, the 120 overlapping genes were highly enriched for genes involved in ER stress-relevant biological processes, such as posttranslational protein targeting to the membrane (p< 2.7×10^-8^) **(Figure 7b)**.

**Figure 7.**
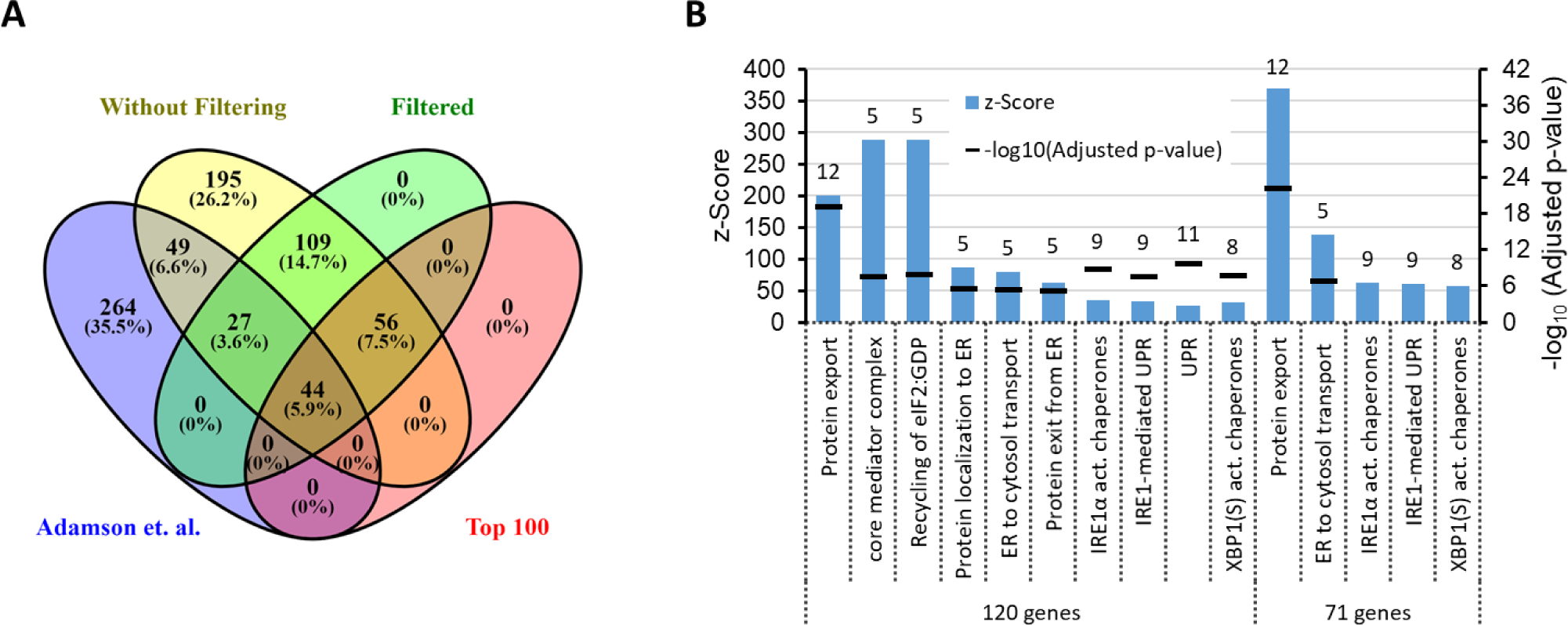
The Gene Signature Module identifies genes that could regulate a given gene signature. a. A Venn diagram representing the intersection of genes implicated in inducing ER stress, as was previously identified through a CRISPRi screen (7), and those detected using the Gene Signature Module. The Gene Signature Module employed the following gene signature: HSPA5+DDIT3+EDEM1+PPP1R15A+HERPUD1+DNAJC3+DNAJB9+DNAJB11+MANF +HERPUD1+SDF2L1+HSP90B1+SELENOK+CDK2AP2+CALR-TUBB-SLC25A3-PTMA-PRDX1-PPIA-TUBB4B-HSPE1-CD59. The label ‘Filtered’ represents the remaining hits after the activation of the filter for blacklisted perturbations (employing a z-score threshold of 2). ‘Top100’ denotes the top 100 hits. **b.** Results of the GSEA conducted for the 120 and 71 genes that overlapped between the previous CRISPRi study and the genes identified by the Gene Signature Module, with and without the application of a filter, respectively.

As it was discussed above in the “Gene Expression Analysis Module” subsection of Methods, certain perturbations have the capacity to elevate global mRNA levels, thereby giving rise to false-positive results in gene signature analyses. Activation of the noise filter for black-listed sgRNAs resulted in the elimination of more than half of the initial hits, effectively reducing the total number of identified genes to 236. Consequently, the overlap with the aforementioned set of 397 genes decreased to 71 **(Figure 7a)**. A major portion of these 71 genes, 44 in total, were included among the top 100 genes identified by our gene signature. These 71 genes were also highly enriched for ER stress relevant processes **(Figure 7b)**. The 264 genes that were exclusively identified in the CRISPRi study were likely inducing ATF4 signaling, as this list included more than 60 mitochondrial proteins, including >35 mitochondrial ribosomal proteins. Targeting these genes may induce ATF4 signaling through the mitochondrial stress response pathway, but not the unfolded protein response (37). Consistently, according to the GO-BP database, among the 264 genes only three genes were implicated in the unfolded protein response. These findings highlight the potential of the Gene Signature Analysis Module in efficiently filtering out noise and extracting key biological insights from GWPS data.

## DISCUSSION

The Perturb-Seq technique is an invaluable tool for probing a wide range of biological processes and gene-function relationships (7–11,15). As the technology continues to advance, it is highly likely that Perturb-Seq will continue to be refined, providing even greater insights into the molecular and cellular mechanisms that underlie these processes. In addition, with the cost of next-generation sequencing continuing to decrease, there will be additional GWPS studies in the near future in many other cell lines and tissues and under different conditions. Therefore, it is important to be prepared for this new era of genomics research and develop platforms that allow users to easily access and query the Perturb-Seq data for their specific needs.

The GeneSetR project, the first phase of which we present here, is currently under active development, and we will introduce several new technical features and capabilities in the near future. For instance, we are currently using the Apache eCharts library for visualizing generated networks. Although visually appealing, this library is not ideal for drawing networks when the number of nodes is high. Therefore, we intend to incorporate the cytoscape/js or vis.js libraries to provide users more options to visualize analysis results that would be more favorable for large networks.

Moreover, in the future, we will enhance GeneSetR by incorporating annotation from other public datasets. For instance, we will incorporate correlation data from the TCGA dataset to generated networks, and add the ability to filter genes based on their expression in normal and cancer tissues. Importantly, we plan to build an active developer community to further expand the functionalities of GeneSetR; we thus encourage all interested parties to participate in the development process via our Github page.

## Data availability

GeneSetR can be freely accessed at https://genesetr.uio.no web site. The source code is hosted on Github (https://github.com/omerfaruk84/GenesetR).

## Funding

This work supported by a grant from the Norwegian Research Council (#303353) and Norwegian Cancer Society (#247110) to FS.

## Acknowledgements

We wish to express our gratitude to Jotun Hein, Ralph Skotheim, Eivind Høvig, Marieke Kuijjer, and Ladislav Hovan for their constructive feedback and insights during the development of GeneSetR and the preparation of the manuscript. Additionally, our thanks go to Klevin Bazaiti for his significant contributions to the front-end development of GeneSetR.

## Notes

https://genesetr.uio.no/

### Competing Interest Statement

The authors have declared no competing interest.

