## Supplementary Information and Figures for "GeneSetR: A web server for gene set analysis based on genome-wide Perturb-Seq data"

**Noise Filters in GeneSetR**

RNA-Seq experiments can be affected by two main types of noise that the Gene Expression Analysis Module aims to filter out. The first type of noise pertains to the perturbations that affect global mRNA levels. To detect these, for each perturbation we computed the average z-scores of all detected genes. As anticipated, the data exhibited a normal distribution around zero, with several outlier values, particularly on the positive side **(Supp. Figure 1a)**.

We identified and filtered out these outliers based on a threshold value of 3.5 modified z-score, following previously established guidelines on outlier detection (38). Subsequently, we calculated average and standard deviation values from the filtered data and used these for regular z-score calculation. The z-score distribution of the results showed that at 2.58 z-score cutoff, which corresponds to 99% Confidence Interval (CI), knockdown of 116 genes led to a global decrease in gene expression **(Supp. Figure 1a** and **Supp. Table 3)**. These genes were enriched for processes such as mitochondrial protein translation, mitochondrial tRNA aminoacylation, histone acetyl transferase activity, mTORC1 mediated signaling, and mitochondrial electron transport, all of which would be expected to affect global gene expression based on their known functions **(Supp. Figure 1b)**.

Among the top 16 hits, three (KAT8, KANSL2, KANSL3) were members of the non-specific lethal (NSL) complex, a chromatin-modifying complex involved in regulation of gene expression with roles in mitochondria and cell division, among others (39). The top hit, KAT8, is the catalytic subunit of the NSL complex, a histone acetyltransferase that acetylates histone H4 at lysine 16 (H4K16ac), a modification associated with activation of transcription (39). Second top hit, SIRT7, is a NAD-dependent protein-lysine deacetylase, which promotes RNA transcription mediated by polymerase I and II (40,41). These results are logical, but the >2.58 (99% CI) z-score cutoff can be considered stringent, as when it is relaxed to >1,96 (95% CI) the list gets expanded to 231 genes, and the additional 115 genes were enriched in similar processes as above.

On the other side of the distribution, at 2.58 (99% CI) z-score cutoff, knockdown of 636 genes resulted in a global increase in gene expression **(Supp. Figure 1a** and **Supp. Table 3)**. This list was predominantly enriched for genes involved in ribosome biogenesis ($p<4.5 \times 10^{-199}$), transcription from RNA polymerase II promoter ($p<3.3 \times 10^{-42}$), and regulation of transcription ($p<0.009$) **(Supp. Figure 1b)**. Top 20 genes included 12 ribosomal proteins, suggesting that interference with ribosomal translation could result in increased global mRNA levels through upregulation of transcription as a compensatory mechanism, or through slowdown of mRNA degradation (42). Not surprisingly, 10 of the 11 members of the exosome complex, which is responsible for the 3'-5' degradation of RNA molecules, were among the top hits.

In addition to perturbations that affect global mRNA synthesis, the second type of noise in RNA-Seq experiments arises from abnormal expression of certain genes in response to perturbations. In general, the expression of a gene is expected to be up- or down-regulated by a comparable number of perturbations, and the average of the z-scores should form a normal distribution around zero. Evaluating whether the expression of specific genes exhibits an unusual distribution compared to others can help identify this type of noise. To address this, we capped the absolute expression values at 1.5 (e.g., values greater than 1.5 was replaced with 1.5 and values less than -1.5 was replaced with -1.5) to mitigate the effects of outlier values. Following this approach, we computed z-scores using average and standard deviation values calculated after outlier removal using modified z-score transformation.

At 99% CI, 157 and 112 genes showed aberrant down- and up-regulation profiles, respectively **(Supp. Figure 1c** and **Supp. Table 4)**. PRG2 was the most frequently upregulated gene and was induced by 933 perturbations; interestingly, no single perturbation down-regulated it. Conversely, the most frequently down-regulated gene was PTMA, which was down-regulated by 717 perturbations and up-regulated by only 13 perturbations.

While it is plausible that certain 'response genes' remain mostly silent until a stimulus triggers their activation, hence they are more likely to be upregulated by perturbations, it is not immediately clear what drives these abnormal expression profiles for most of the identified genes. GSEA suggested that 'iron homeostasis' and 'mRNA splicing' related genes were enriched among the genes that tend to be up- or down-regulated, respectively **(Supp. Figure 1d)**. However, this could merely be linked to the high expression of the genes involved in these processes. Enrichment analysis using the 'Enrichr Libraries Most Popular Genes' library suggested that several of these genes were frequently identified as deregulated upon various genetic or chemical perturbations in RNA-Seq experiments, implying that these genes could be introducing noise in these analyses due to specific biological properties (e.g., high expression, high variability in expression, etc.) or due to technical biases present in RNA-sequencing (more abundant RNAs being represented higher). A deeper investigation is needed to untangle the molecular or technical mechanisms behind these patterns.

**Benchmarking of dimensionality reduction algorithms**

Since the number of components (NOC) is a key setting for all DR algorithms, in order to systematically evaluate the optimal value for it, and to determine which DR algorithm would result in functionally relevant clusters, we conducted a benchmarking study using four DR algorithms available in GeneSetR, as well as some of their combinations. Using a script, we varied the NOC value and assessed the number of generated clusters, the total number of genes included in the generated clusters, and the computational time spent during the calculations. Moreover, in the script, to assess whether the generated clusters are functionally relevant, GSEA was performed in each cluster using the GO-BP database, and the number of genes in the top enriched term was used to calculate a “cluster score” (square of number of genes in top enriched term / number of genes in the cluster) **(Supp. Figure 2a)**. The sum of the scores of all generated clusters for the selected NOC value was used as the “clustering efficiency score” for that particular NOC setting.

For the PCA we ran the benchmarking script with varying values of explained variance (EV) between 30 and 100 percent **(Supp. Figure 2b-c)**. As expected, PCA-reduced data did not yield many clusters and only two to five clusters were formed for varying levels of EV. The number of clusters and the total number of clustered genes tended to decrease as the EV values increased. Although three to four clusters were obtained at low EV values (<45%), their score were quite low. The clustering score reached its maximum value of 35 at 60% EV, after which it gradually decreased with increasing

EV values. At the peak clustering score, approximately 100 genes were included in the clusters.

For the UMAP, we varied the NOC value between three and 398 (which corresponds to the size of the available sgRNAs for the gene list under consideration) **(Supp. Figure 2d-e)**. The results revealed that both the number of generated clusters and clustering score exhibited a certain degree of variability across the range of examined NOC values. However, we did not observe any discernible trend in either of these metrics with respect to the NOC. Consistently, the total number of clustered genes remained relatively stable, hovering around 280 across all tested component settings. At the peak clustering score of 44, a total of 12 clusters were formed. These findings suggest that the choice of the number of components may not have a significant impact on the overall clustering performance of the UMAP algorithm, at least within the range of values tested in our study.

On the other hand, for tSNE, we found that in addition to the NOC setting, the algorithm exhibits significant sensitivity to the perplexity parameter, which determines the balance between preserving the local and global structure of data by setting the effective number of nearest neighbors. According to the tSNE user manual, the suggested perplexity default value is 30, and it is advised to keep it between 5 and 50 in most cases. However, when running our benchmark script with that setting, it resulted in very low clustering scores (<11) for varying NOC values **(Supp. Figure 3a-b)**. The best score of 11.2 was obtained at NOC value of three and then it sharply decreased to values less than 2. Therefore, we investigated the effect of varying perplexity values from 1 to 300 while maintaining a constant NOC value of three.

tSNE was capable of producing 19 distinct clusters with a total clustering score of 47 at the lowest perplexity value **(Supp. Figure 2f-g)**. As the perplexity value was increased, the number of identified clusters decreased dramatically to a few clusters. However, a sudden increase in clustering efficiency was observed around the perplexity value of 150, albeit the clustering score and number of clusters that are still lower compared to those obtained with a perplexity value of one. Moreover, increasing the NOC value to 10 while maintaining the perplexity value of one greatly impeded the clustering process. Therefore, to find optimal settings, we conducted a more comprehensive analysis by varying both perplexity and NOC values and generated heatmaps for clustering efficiency and the number of obtained clusters. However, it was not possible to get a better clustering score in our test range **(Supp. Figure 3c)**.

On the other hand, MDE algorithm was unsuccessful in handling high NOC values, thus limiting the ability to test the algorithm beyond the range of 3 to 163. Notably, the highest clustering score of 48 was attained when the NOC value was set to 43 **(Supp. Figure 2h)**. Prior to this point, a modest upward trend in clustering scores was observed, followed by a relatively sharp decline beyond this threshold. Concurrently, the total number of clustered genes exhibited a gradual decline as the NOC values increased **(Supp. Figure 2i)**. At the optimal clustering score, the HDB-Scan algorithm was able to identify up to 15 distinct clusters from the MDE-reduced data.

We then investigated whether a sequential application of DR algorithms would yield improved clustering of the data. To this end, we first employed PCA with a 60% explained variance setting, resulting in 27 components, and subsequently applied the remaining non-linear DR algorithms on the PCA-reduced data. We evaluated the clustering efficiency for the NOC values ranging from three to 27. Surprisingly, UMAP yielded a maximum clustering score of 55 at the lowest NOC value, resulting in 14 clusters comprising approximately 340 genes **(Supp. Figure 4a-b)**. This was a significant improvement compared to the UMAP-only run (CS: 44, 12 clusters). For higher NOC values, the clustering score fluctuated around 48, with a corresponding cluster count of approximately 9, demonstrating a slightly decreasing trend. When tSNE was applied, a maximum clustering score of 36 was obtained at the NOC value of 10, resulting in 9 clusters comprising 134 genes **(Supp. Figure 4c-d)**. While keeping the NOC value at 10, varying the perplexity value between 1 and 50 showed that at the perplexity value of 7, slightly better clustering results (clustering score ~48) can be achieved **(Supp. Figure 4e-f)**. Finally, the combination of PCA with MDE yielded the best clustering scores in the sequential analysis. With increasing NOC values of up to 19, the clustering scores gradually increased up to 80 **(Supp. Figure 4g-h)**. At this peak level, 22-23 clusters were generated which covered almost all genes. 12 of the 23 clusters had a cluster score greater than one and the top enriched terms for these clusters had p values less than $1x10^{-7}$ **(Supp. Table 5)**.

During the course of these optimization studies, we also monitored the CPU time utilized by each DR algorithm. The computational cost of the PCA was independent of the number of output dimensions, with average run times of less than 400 milliseconds (using an i7-7700 CPU @3.6 GHz, 32 GB RAM computer running Windows 10) **(Supp. Figure 5)**. In contrast, the run times of the three non-linear DR algorithms - UMAP, tSNE and MDE- exhibited varying degrees of correlation with the number of output dimensions. Specifically, UMAP and MDE showed a linear correlation, with UMAP requiring a minimum of around five seconds for outputting three dimensions and gradually increasing to approximately eight seconds for outputting 393 components. While MDE necessitated only one second for outputting three dimensions, its computational cost rose more steeply than that of UMAP, until ∼160 components, beyond which it failed to execute successfully. Of the four DR algorithms studied, tSNE was the most computationally expensive one requiring three seconds to output three dimensions, but experiencing a dramatic increase to almost five minutes for outputting 393 components. Notably, while the cost increase was generally linear across most intervals, at certain points there were sharp increases. For instance, the calculation of 323 components required 109 seconds, whereas the calculation of 328 components required more than twice that amount, costing 242 seconds of CPU time. In summary, considering the clustering efficiencies and runtimes, a combination of PCA and MDE produced the most efficient clustering results.

**Supp. Figure 1. Identification of noise in the Perturb-Seq data**
**a.** Histogram in the middle shows distribution of average effect of each perturbation across all detected genes. The left and right plots show perturbations that lead to a general decrease or increase in total mRNA levels, respectively. **b.** Shows the results of the GSEA of the significant perturbations (z-score >1.95) highlighted in **a**. **c.** Histogram shows distribution of average expression level of each gene across all perturbations. The left and right plots show genes that are commonly up- or down-regulated by perturbations, respectively. **d.** Shows the results of the GSEA of the significant genes (z-score >1.95) that were highlighted in **c**. In **a** and **c,** functionally related genes (e.g., those that are members of the same family, or form a functional complex), are highlighted in same color. Note that as shown in **b,** interfering with mitochondrial translation or histone acetylation negatively affects total mRNA expression, whereas inhibition of processes such as ribosome biogenesis and mRNA catabolism positively affects total mRNA expression. Also note that, as shown in **d**, genes involved in certain biological processes, such as mRNA splicing and ribosome biogenesis, tend to be down-regulated, whereas genes involved in iron homeostasis and regulation of NF-κB signaling tend to be upregulated upon perturbation of various genes. In addition, genes that tend to be down-regulated or up-regulated were enriched in single gene, drug or disease perturbation datasets harboring down- or up-regulated genes, respectively, suggesting that these genes are identified due to certain biological properties or technical reasons.

**Supp. Figure 2. Benchmarking of dimensionality reduction algorithms. a.** A cluster score is calculated through multiplying the number of genes in top enriched term with its ratio to the cluster size (CS). Therefore, the score can vary between 1/CS and CS and responds both to the cluster size and the number of overlapping genes with top enriched biological process. If the overlap is small, the score drops significantly (i); if the overlap is high, depending on the size of the cluster, score increases (ii and iii). Clustering efficiency score for a particular dimensionality reduction run is calculated as a sum of cluster scores for the generated clusters (right panel). **b-i.** Line plots are presented for benchmarking results for the indicated dimensionality reduction algorithms. Error bars show SEM for 10 iterations.

**Supp. Figure 3. Benchmarking of the t-SNE algorithm. a** and **b.** Line plots represent benchmarking results for the t-SNE algorithm executed with increasing values for the number of components. **c.** A 2D heatmap illustrates clustering scores for varying values of the number of components and perplexity settings of the t-SNE algorithm.

**Supp. Figure 4. Benchmarking sequential execution of dimensionality reduction algorithms. a-f.** Line plots are presented from benchmarking results for the indicated sequentially run dimensionality reduction algorithms. Error bars show SEM for 10 iterations.

**Supp. Figure 5. Comparison of computation times for dimensionality reduction algorithms.** Line graph displays the mean computational run time taken by the specified algorithms. Error bars show SEM for 10 iterations.

**Supp. Table 1**: Simplified list of interactions identified by Gene Expression Analysis module for ATF4 gene using the settings below. Z-score threshold: 0.4 correlative links: Excluded Black listed sgRNA threshold: 2 (Directional Only) Black listed genes threshold: 2 (Directional Only) Perturbation Count Filter: 500 Gene Expression Count Filter: 500. NC stands for neighbor count. Score stands for Z score. Positive z Scores indicate upregulation. Negative Z scores indicate downregulation. UNR: Upstream Negative Regulator (so a perturbation that cause upregulation of GOI). UPR: Upstream Positive Regulator (so a perturbation that cause downregulation of GOI). DNR: Downstream Negatively Regulated (so a gene that is upregulated upon perturbation of GOI). DPR: Downstream Positively Regulated (so a gene that is downregulated upon perturbation of GOI). UNR_DNR: A link between an upstream negative regulator (source) and a downstream negatively regulated.

**Supp. Table 2:** Full list of interactions identified by Gene Expression Analysis module for ATF4 gene using the settings below. Z-score threshold: 0.4 correlative links: Excluded Black listed sgRNA threshold: 2 (Directional Only) Black listed genes threshold: 2 (Directional Only) Perturbation Count Filter: 500 Gene Expression Count Filter: 500. NC stands for neighbor count. Score stands for Z score. Positive z Scores indicate upregulation. Negative Z scores indicate downregulation. UNR: Upstream Negative Regulator (so a perturbation that cause upregulation of GOI). UPR: Upstream Positive Regulator (so a perturbation that cause downregulation of GOI). DNR: Downstream Negatively Regulated (so a gene that is upregulated upon perturbation of GOI). DPR: Downstream Positively Regulated (so a gene that is downregulated upon perturbation of GOI). UNR_DNR: A link between an upstream negative regulator (source) and a downstream negatively regulated.

**Supp. Table 3:** Z-scores of average effect of each perturbation across all detected genes. Perturbations that have absolute z-Score value greater 2.58 (99% confidence interval) are colored in red, while the ones that are between 2 and 1.96 (95% confidence interval) are colored in light blue. Up and Down columns shows the number of genes that are affected from the particular perturbation based on 0.4 threshold.

**Supp. Table 4:** Z-scores of average expression level of each gene across all perturbations. Genes that have absolute z-Score value greater 2.58 (99% confidence interval) are colored in red, while the ones that are between 2 and 1.96 (95% confidence interval) are colored in light blue. Up and Down columns show the number of perturbations that affected expression of the particular gene based on 0.4 threshold.

**Supp. Table 5:** The table presents the outcomes of the gene set enrichment analysis and cluster scores for the clusters generated by the HDBSCAN algorithm (minimum cluster size: 5; cluster selection epsilon: 0; and metric: Euclidean) following dimensionality reduction through a sequential application of PCA (60% explained variance) and MDE (number of components: 21) algorithms.

### Supplementary Figure 1 – Identification of noise in the perturb-seq data

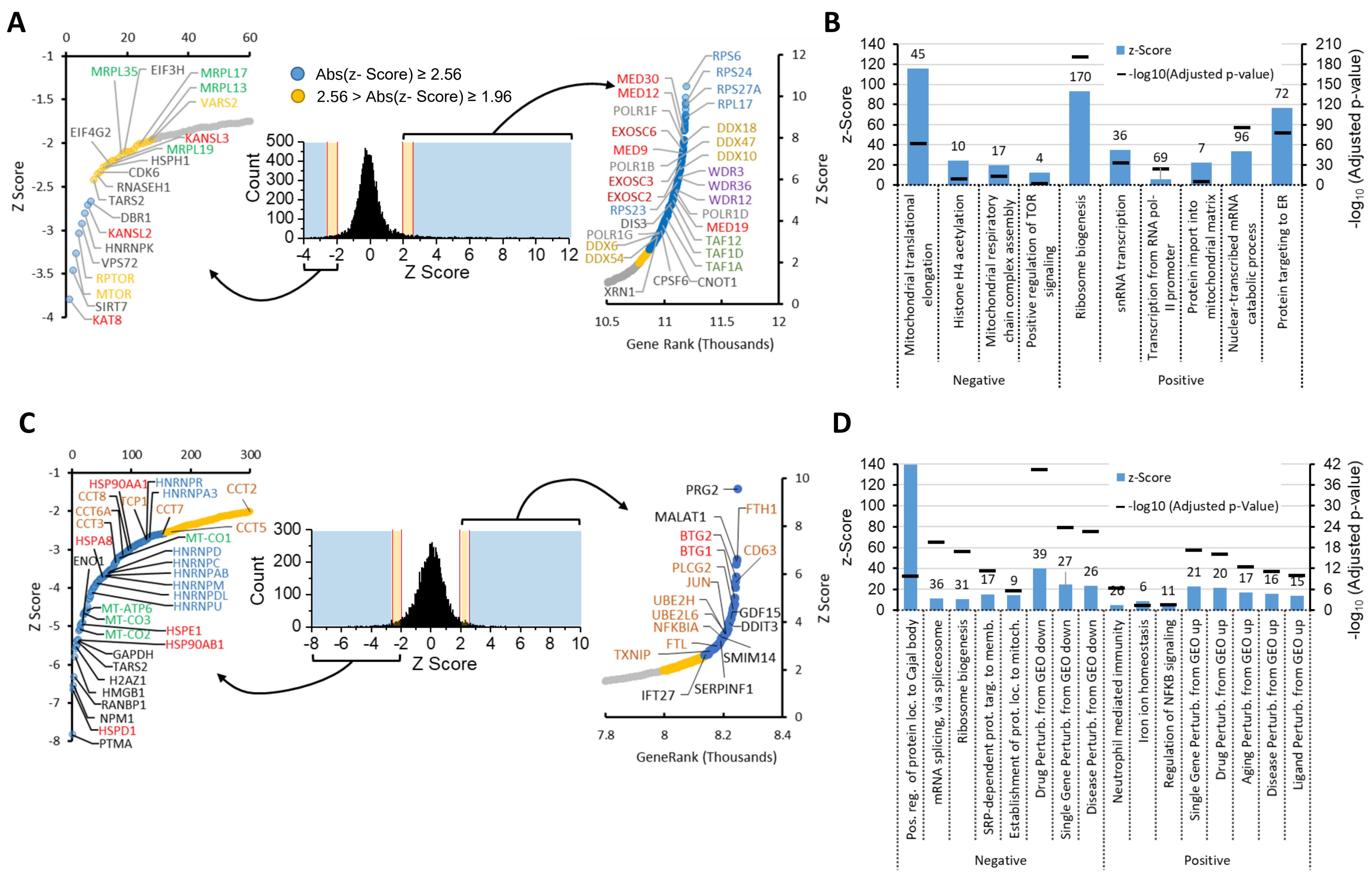

### Supplementary Figure 2 – Benchmarking of dimensionality reduction algorithms

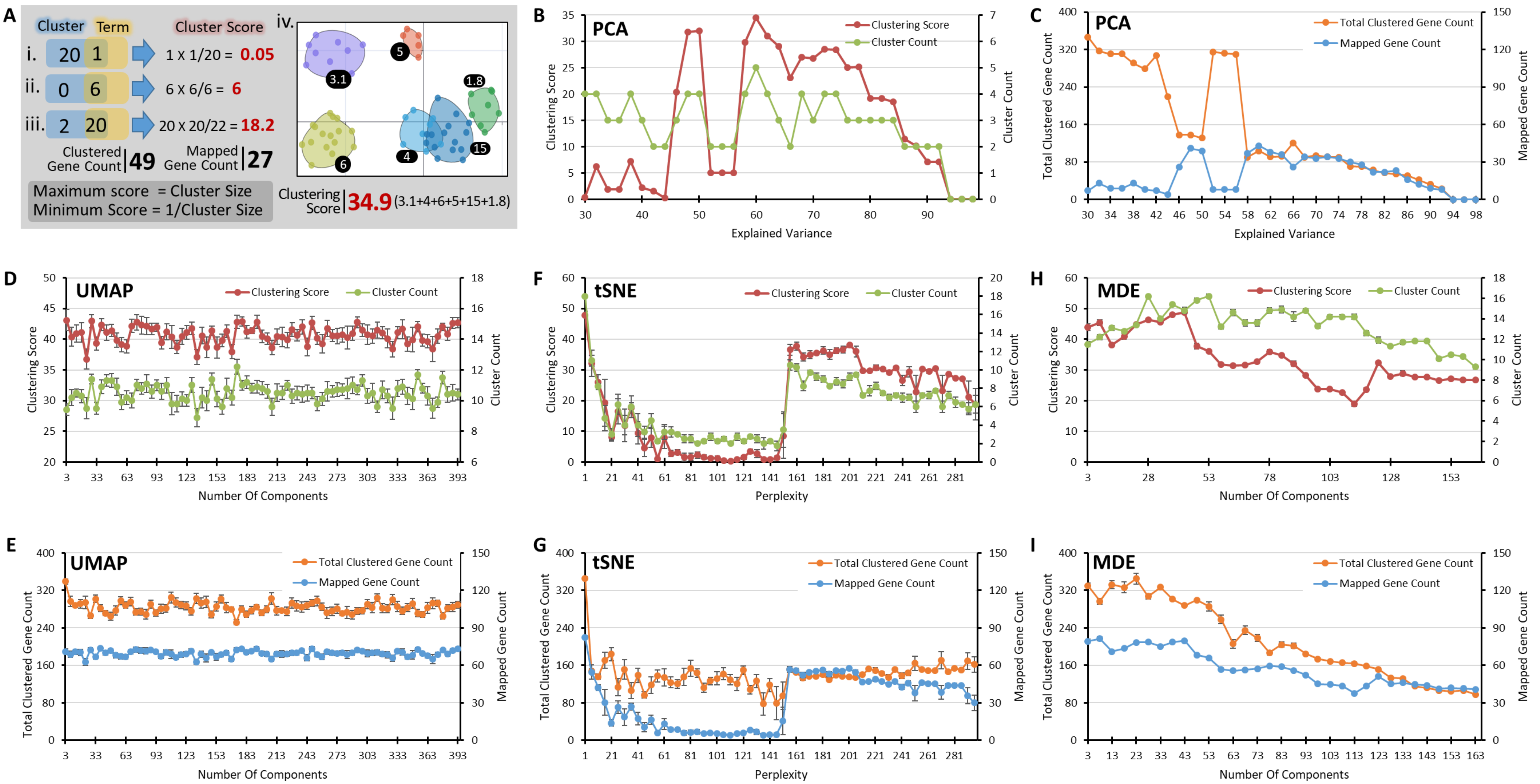

Supplementary Figure 3 – Benchmarking of the tSNE algorithm

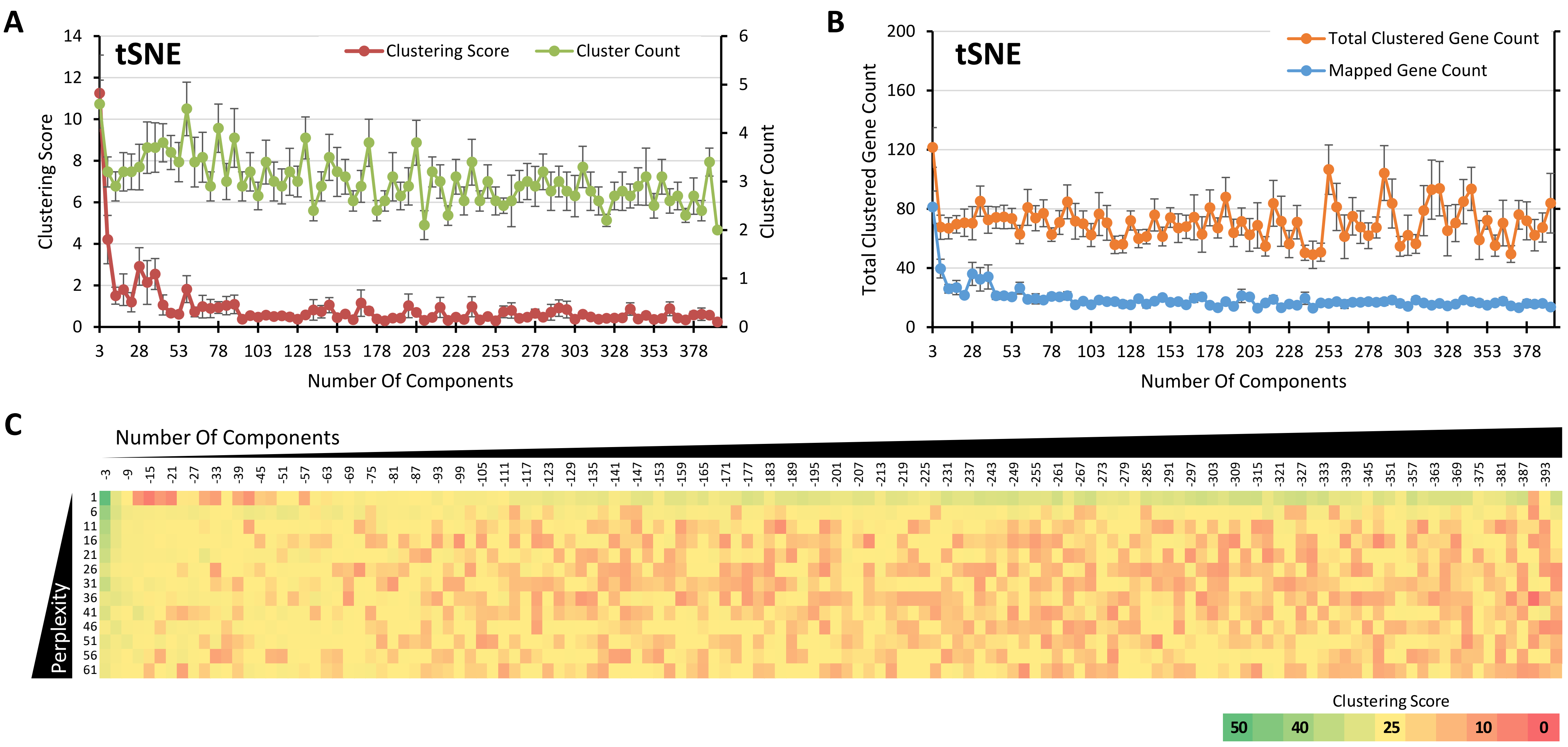

### Supplementary Figure 4 – Benchmarking of dimensionality reduction algorithms executed sequentially

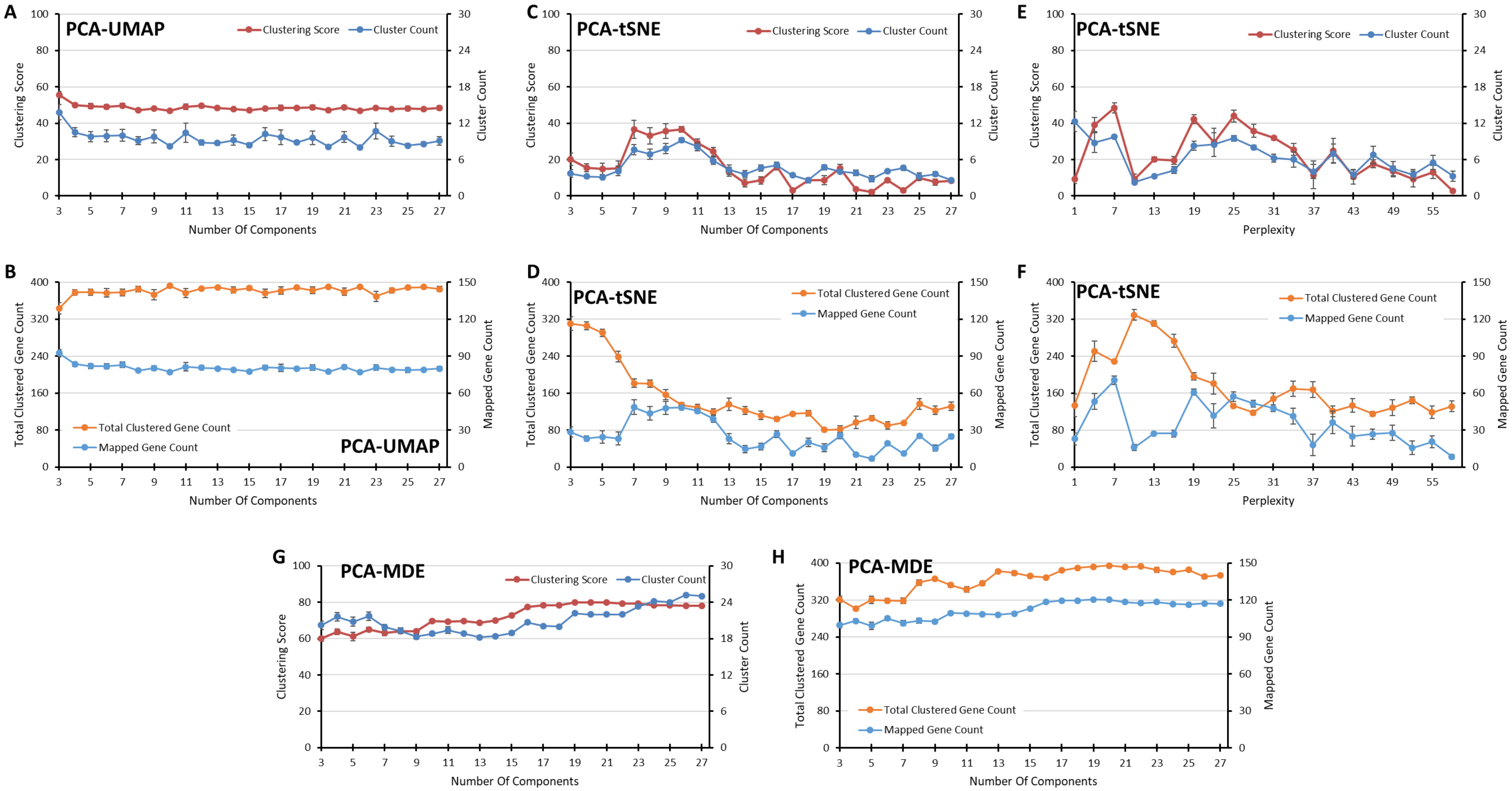

Supplementary Figure 5 – Comparison of Efficiency and Calculation Time for Various DR Algorithms

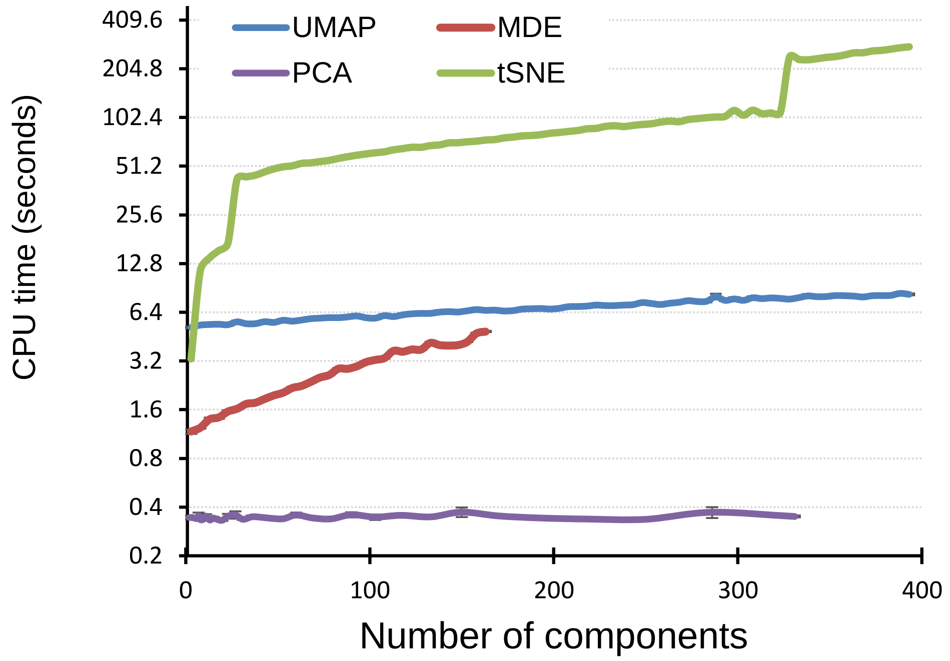
